## Supplementary Material for "Functionnectome: a framework to analyse the contribution of brain circuits to fMRI"

**Region-wise analysis**

*Principle*

The Functionnectome has the ability to project the functional signal of each brain voxel onto the white matter. Although the principle is relatively straightforward, the computation can require up to several hundred thousand steps per subject, depending on how many voxels are kept for the analysis. This is quite computationally intensive and thus requires powerful computers and servers, which are not necessarily available to all. In order to make the Functionnectome accessible to the broadest audience, we added the possibility to use brain regions instead of voxels as spatial units, reducing the number of processing steps to a few hundred per subject. This simplification of the analyses decreases the computation time from hours per subject (or even days on consumer-grade computers) to minutes. The brain regions used for this example of simplified analysis, and the associated probability maps, were derived from the HCP multi-modal parcellation (MMP) and are included in the priors provided with the software. During the analysis, the functional signal from a given region at a specific time-point is defined as the median of the values of that region’s voxels at that time-point. Here, we used the median instead of the mean in order to mitigate the effect of noisy voxels in small regions.

*Results*

All the analyses presented in the main body of the article were replicated using the region-wise analysis aforementioned. For the sake of comparison, we kept the same activation-values threshold as in the original analyses presented in the main text.

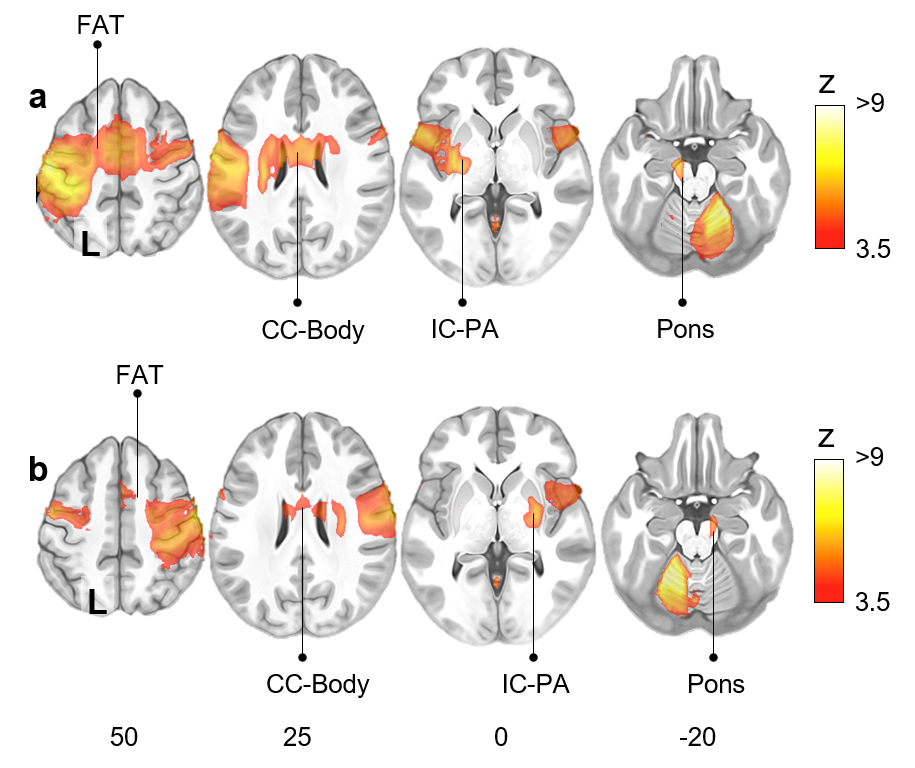

Supplementary figure 1: Right (a) and left (b) finger-tapping motor activation network from the region-wise Functionnectome analysis. FAT: Frontal Aslant Tract; CC-Body: Corpus Callosum body; IC-PA: Internal capsule posterior arm.

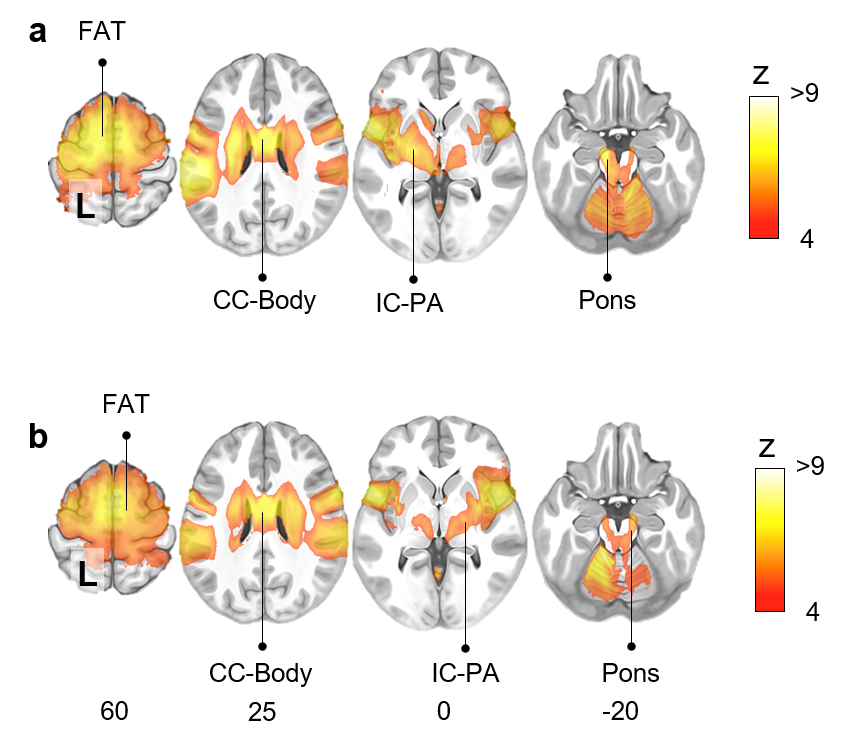

Supplementary figure 2: Motor task activation network for left (a) and right (b) toes clenching, from the region-wise Functionnectome analysis. FAT: Frontal Aslant Tract; CC-Body: Corpus Callosum body; Pons; IC-PA: Internal capsule posterior arm.

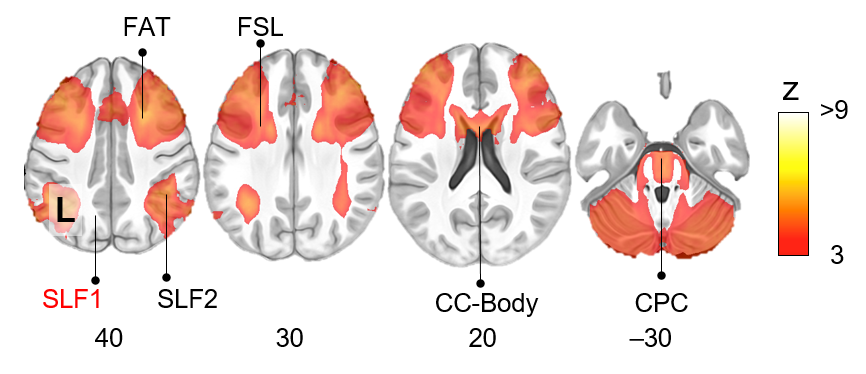

Supplementary figure 3: Working memory task activation network from the region-wise Functionnectome analysis. Pathways not present here but appearing in the voxel-wise analysis are labelled in red. FAT: Frontal Aslant Tract; FSL: Frontal Superior Longitudinal tract; SLF: Superior Longitudinal Fasciculus; CC-Body: Corpus Callosum body. CPC: Cortico-Ponto-Cerebellar tract. Tracts indicated in red were identified in the original voxel-wise analysis but missing in the simplified region-wise analysis.

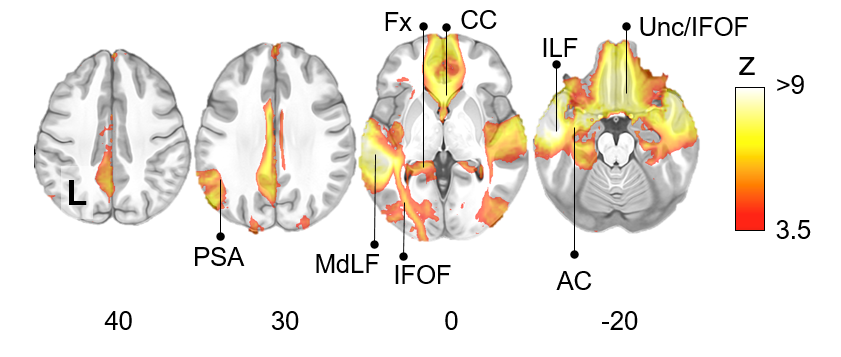

Supplementary figure 4: Semantic system task activation network from the region-wise Functionnectome analysis. Pathways appearing in the voxel-wise analysis but not present here are labelled in red. PSA: Posterior Segment of the arcuate fasciculus; MdLF: Middle Longitudinal Fasciculus; IFOF: Inferior Fronto-Occipital Fasciculus; Fx: Fornix; CC: Corpus Callosum; ILF: Inferior Longitudinal Fasciculus; Unc: Uncinate fasciculus; AC: Anterior Commissure.

*Supplementary discussion*

The region-wise analysis delivers results comparable to those obtained with the voxel-wise analysis, albeit slightly less sensitive and specific. For instance, compared to the voxel-wise analysis, a few pathways were missing in the region-wise analysis (e.g. the superior longitudinal fasciculus 1 in the working memory task map - Supplementary figure 3), and other tracts were displaying a larger activations (e.g. the first branch of the superior longitudinal fasciculus - SLF1 - in the working memory task - Supplementary figure 3). The choice of brain segmentation used for the simplified region-wise analysis could improve the results. Here, the multimodal parcellation of the HCP has a general-purpose, but using a brain parcellation tailored for the specific analysis of a project could yield better results.

Overall, the region-wise analysis offers a way for any research team to use the Functionnectome and obtain results in a short amount of time, independently from the computational resources available. Even teams considering the application of the voxel-wise analysis could use it as a quick test of their method, before investing the time and computational resources for the complete analysis.
